# A human cytomegalovirus prefusion-like glycoprotein B subunit vaccine elicits similar humoral immunity to that of postfusion gB in mice

**DOI:** 10.1101/2024.11.18.624140

**Authors:** Krithika P. Karthigeyan, Megan Connors, Christian R. Binuya, Mackensie Gross, Adelaide S. Fuller, Chelsea M. Crooks, Hsuan-Yuan Wang, Madeline R. Sponholtz, Patrick O. Byrne, Savannah Herbek, Caroline Andy, Linda M. Gerber, John D. Campbell, Caitlin A. Williams, Itzayana Miller, Dong Yu, Matthew J. Bottomley, Jason S. McLellan, Sallie R. Permar

## Abstract

Human cytomegalovirus (HCMV) is the leading infectious cause of birth defects. Despite the global disease burden, there is no FDA-approved HCMV vaccine. The most efficacious HCMV vaccine candidates to date have used glycoprotein B (gB), a class III viral fusion protein, in its postfusion form. While some viral fusion proteins have been shown to elicit stronger neutralizing responses in their prefusion conformation, HCMV prefusion-like and postfusion gB were recently shown to elicit antibodies with similar fibroblast neutralization titers in mice. We aimed to define and compare the specificity and functionality of plasma IgG elicited by distinct prefusion-like and postfusion gB constructs. Prefusion-like and postfusion gB elicited comparable IgG responses that predominantly mapped to AD-5 antigenic domain known to elicit neutralizing antibodies. Interestingly, postfusion gB elicited significantly higher plasma IgG binding to cell-associated gB and antibody dependent cellular phagocytosis than that of prefusion-like gB. The vaccines elicited comparable neutralization titers of heterologous HCMV strain AD169r in fibroblasts, yet neither elicited neutralizing titers against the vaccine-matched strain Towne in fibroblasts. Our data indicate that gB in this prefusion-like conformation elicits similar specificity and functional humoral immunity to that of postfusion gB, unlike certain class I viral fusion proteins which have been used as vaccine antigens. These findings deepen our understanding of the immune response elicited by class III fusion proteins and may inform further design and testing of conformationally-dependent herpesvirus glycoprotein vaccine candidates.

## INTRODUCTION

Despite substantial efforts over five decades, there is still no licensed vaccine against the leading infectious cause of birth defects worldwide, human cytomegalovirus (HCMV)^1^. HCMV infection is asymptomatic in healthy adults, but can cause devastating sequelae in immunocompromised subjects, including organ transplant recipients^2^ and children who acquired the infection *in utero* or postnatally as pre-term infants^3^. The burden of congenital HCMV is highest in Black and Hispanic populations^4^, in low and middle-income countries,^5,6^, and in pre-term infants^7,8^. In the US, about 1 out of 3 pregnant people primarily infected with HCMV during pregnancy will pass the virus to the fetus, which accounts for nearly 1 in 200 children born with congenital HCMV^9^. At 235 kbp, HCMV has one of the largest genomes of any human virus. The HCMV genome exhibits high levels of genetic diversity and encodes multiple glycoproteins that are involved in entry and fusion into fibroblast, endothelial, and epithelial cells^10,11,12^. While the fusion protein glycoprotein B (gB) has been the primary target of HCMV vaccine efforts^13–15^, the pentameric complex (PC) composed of gH, gL, UL128, UL130, and UL131A has also been investigated in recent trials^16^, as well as certain targets of cellular immunity, including tegument protein pp65 and the immediate early proteins 1 and 2 (IE1, IE2)^17^.

As a result of long-term co-evolution and adaptation, HCMV is a highly species-specific virus that has evolved complex immune evasion mechanisms^18^, which make vaccine efforts challenging. The most successful vaccine candidate to date is a subunit vaccine that contains glycoprotein B (Sanofi) formulated with the oil-in-water emulsion adjuvant MF59 (Novartis). The gB/MF59 vaccine was administered as a three-dose series to seronegative adolescent and postpartum women in different trials and achieved nearly 50% efficacy, albeit short-lived, in multiple phase 2b clinical trials^14,15^. While this vaccine elicited poorly neutralizing antibody responses, plasma IgG binding to gB expressed on the surface of a cell (cell-associated gB) was identified as a correlate of protection for this vaccine^19^. Other studies have also implicated non-neutralizing antibody functions in the observed clinical efficacy of the gB/MF59 vaccine, such as antibody-dependent cellular phagocytosis (ADCP)^20^ and prevention of cell-cell spread of HCMV^21^. Though the HCMV vaccine field has been investigating other glycoprotein antigens in addition to gB and/or novel vaccine platforms as vaccine candidates^22^, interest in a subunit gB vaccine— particularly one modified to display the gB antigen in its prefusion conformation—has been reinvigorated by the recent determination of the prefusion structure of HCMV gB and the successes of the RSV and SARS-CoV-2 vaccines, which present the class I viral fusion proteins in their prefusion state^23–25^.

HCMV gB is a homotrimeric class III fusion protein that brings the viral and host cell membranes together when it transitions from a metastable prefusion state to a highly stable postfusion state, thereby facilitating viral facilitates host cell entry^26^. The ectodomain of gB is composed of five structural domains (I – V) which, together with the membrane proximal region, the transmembrane domain, and the C-terminal domain, are classified functionally into six antigenic domains (AD 1-6)^21,27^. The immunodominant AD-1 domain on structural domain IV is the target of mostly non-neutralizing antibodies.^28^ AD-2 corresponds to the first 85 residues of gB and can be further classified into AD-2 site 1, the target of potently neutralizing antibodies, and AD-2 site 2, which elicits exclusively non-neutralizing antibodies. AD-3 on the C-terminal domain and AD-6 on domain V also elicit exclusively non-neutralizing antibodies.^21,29,30^ On the other hand, AD-4 and AD-5 on structural domains I and II respectively are targets of neutralizing antibodies^29,31,32^. Antibodies targeting the recently characterized domain AD-6 have been associated with blocking cell-to-cell spread of HCMV^21^.

While many class I fusion proteins have been stabilized in their prefusion conformations, the prefusion conformations of class III fusion proteins— particularly those encoded by herpesviruses— have been more elusive^33^. The first high-resolution published structure of HCMV gB in the prefusion conformation was detergent-solubilized, full-length, membrane-bound gB^34^. This structure was recently used to guide the design of a gB ectodomain in a prefusion-like conformation, termed gB-C7^35^. Though this prefusion-like gB-C7 did not elicit higher fibroblast neutralization titers compared to postfusion gB ectodomain in mice^35^, the characterization of non-neutralizing antibody functions as well as a comprehensive mapping of the binding responses could shed light on a broader profile of humoral immune responses elicited by these two vaccines. Our study immunized mice with prefusion-like or postfusion gB and provides a detailed characterization of the differences in the humoral immunogenicity between prefusion-like and postfusion gB, with attention to the fine specificity of the elicited IgG responses and non-neutralizing antibody functions. Overall, this immunogenicity study informs structure-guided design of HCMV gB vaccine candidates and for class III viral fusion proteins more broadly, while informing HCMV vaccine-elicited immunogenicity assessments.

## METHODS

### Affinity Purification of prefusion-like HCMV gB ectodomain

HEK293F cell cultures (1L) were grown to a density of 1 x 10^6^ cells/mL and then transiently transfected with 0.5 mg plasmid encoding prefusion-like HCMV gB-C7 using PEI. After 6 days, medium was harvested by centrifugation and 0.22 µm filtered. The media was then passed over Strep-Tactin Sepharose resin (IBA Lifesciences), washed with 3 column volumes of Endotoxin-Free Dulbecco′s PBS (Sigma-Aldrich), and eluted with Strep-Tactin elution buffer (100 mM Tris-Cl pH 8.0, 150 mM NaCl, 1 mM EDTA, and 2.5 mM desthiobiotin) (IBA Lifesciences). Elution fractions were analyzed by SDS-PAGE and fractions containing HCMV gB-C7 were pooled. The eluate was then buffer exchanged to Endotoxin-Free Dulbecco′s PBS (Sigma-Aldrich), concentrated with Amicon Ultra centrifugal filters (MilliporeSigma), and flash-frozen in liquid nitrogen.

### Animal Study Design

All animal studies were approved by the Institutional Animal Care and Use Committee (IACUC) of Weill Cornell Medicine under protocol number 2022-0032. 6-8 week old BALB/c mice (Charles River) were immunized with 2 µg of prefusion-like gB-C7 ectodomain (n= 5) or full-length postfusion gB lacking the transmembrane domain (n = 5) (Sino Biologicals) at weeks 0, 3, and 7. Antigens were administered intramuscularly with CpG 1018 adjuvant (10 µg, Dynavax Technologies Corporation) and aluminum hydroxide (50 µg, Invitrogen). Blood was collected weekly by tail vein nick. Immunogenicity analysis was carried out using plasma samples collected two and four weeks post the three-dose series, i.e., at weeks 9 and 11.

### gB Binding Assays

Plasma IgG binding to soluble prefusion-like and postfusion gB was measured by ELISA at a starting dilution of 1:1000 and serially diluted 3-fold (8-point dilution series). 384-well enzyme-linked immunosorbent assay (ELISA) plates were coated overnight at 4°C with postfusion gB (1 µg/well) or prefusion-like gB-C7 (4.5 µg/well) and then blocked in assay diluent (1x phosphate buffered saline (PBS) (pH 7.4) containing 4% whey, 15% normal goat serum, and 0.5% Tween 20). Goat anti-mouse peroxidase-conjugated IgG secondary antibody was used to detect binding. Plates were developed using SureBlue Substrate and optical density was obtained using a Synergy plate reader. . The 50% effective dose (ED_50_) or concentration (EC_50_) was defined as the reciprocal of the sera dilution or concentration of IgG MAbs that caused a 50% effect using non-linear regression analysis (Sigmoidal, 4PL) using GraphPad Prism.

### Cell culture

All cell lines were obtained from the American Type Culture Collection (ATCC). Human retinal pigment epithelial (ARPE-19) cells were maintained in Dulbecco’s modified Eagle medium-12 (DMEM-F12) supplemented with 10% fetal bovine serum (FBS), 50 U/mL penicillin, and 50 μg/mL streptomycin. Human epithelial kidney (HEK293T) cells were maintained in DMEM containing 10% FBS, 25mM HEPES buffer, 50 U/mL penicillin, and 50 μg/mL streptomycin. Human foreskin fibroblast cells (HFF-1) were maintained in DMEM supplemented with 10% FBS, 25mM HEPES buffer, 50 U/mL penicillin, 50 μg/mL streptomycin and gentamicin, and 25 mM L-Glutamine. Human monocyte (THP-1) cells were maintained in RPMI-1640 medium containing 10% FBS. All cell lines were maintained for a maximum of 25 passages and tested for the presence of mycoplasma biannually.

### gB-transfected cell IgG binding

Plasma IgG binding to gB expressed on the cell surface was measured as previously described^19^. Briefly, HEK 293T cells were co-transfected with plasmids expressing GFP and full-length HCMV gB (Towne strain) for 48 hours and then incubated at 37°C with 1:2500 diluted mouse plasma or 1:6250 diluted CYTOGAM^®^ (Source). Cells were stained with Live/Dead Fixable Near-IR Dead Cell Stain, followed by phycoerythrin (PE) - conjugated goat-anti-mouse IgG Fc staining, and fixed with 10% formalin prior to acquisition via high throughput sampler (HTS) on the flow cytometer (Fortessa; BD). The frequency of PE+ cells was reported for each sample based on the live, singlet, GFP+ population. The average + 3 standard deviation (SD) of baseline (Week 0) samples was used as the positivity cutoff.

### HCMV Production

Viruses and Bacterial Artificial Chromosomes (BACs) were a gift from Professor Tom Shenk. Towne virus was propagated on HFF-1 cells in T-175 culture flasks. AD169 revertant virus containing the UL131-UL128 ORF from HCMV TR and expressing GFP (AD169r-BAC-GFP) was propagated from BAC-transfection of HFF-1 cells using Lipofectamine-3000 (Thermo Fisher Scientific) to produce seed stocks. Working stocks were propagated from seed stock infection of ARPE-19 cells in T-175 flasks. Ts15nR (Towne-BAC-GFP with intact pentamer) was propagated in ARPE-19 cells. Supernatant containing cell-free virus was collected when 90% of cells showed cytopathic effects (∼2 weeks), and cleared of cell debris by low-speed centrifugation before ultracentrifugation through a 20% sucrose cushion.

### Neutralization Assays

Plasma samples were heat-inactivated at 56°C for 30 min. Neutralization was measured by immunostaining of HCMV immediate early-1 (IE-1) protein to quantify reductions in virus infection in HFF-1 or ARPE-19 cells for fibroblast and epithelial neutralization respectively. Briefly, 6000 cells were seeded per well of a 384-well plate and incubated at 37 °C overnight. AD169r or Towne (MOI=1) was incubated with plasma IgG (starting dilutions of 1:10 serially diluted 3-fold and incubated) for 1 hour at 37°C in 96-well culture plates. Rabbit complement (Cedarlane) was added at an 8-fold dilution. The anti-HCMV IgG preparation CYTOGAM was used as a positive control^36^.Virus-antibody preparations were incubated with cells for 24-48 hours, followed by fixing and HCMV IE-1 staining and counter-stained with IgG-AF488 and then DAPI. Images were acquired using ImageXpress Pico Automated Cell Imaging System (Molecular Devices). Infected cells are calculated as a percentage of AF488 positive cells relative to total number of cells determined by DAPI staining. The 50% inhibitory dose (ID_50_) or concentration (IC_50_) was defined as the reciprocal of the sera dilution or concentration of IgG MAbs that caused a 50% reduction in infected cells compared to virus control and were calculated on Graphpad Prism using non-linear regression analysis (Sigmoidal, 4PL), with average % virus infection set as the maximum constraint and the minimum constraint set to zero.

### IgG mapping to gB antigenic domains

Vaccine induced IgG binding to HCMV antigens gB AD-1, gB AD-4, gB AD-5, gB AD-4 + AD-5AD4 + AD5 were measured by multiplex assay as previously described^16^. Briefly, antigens were covalently coupled to fluorescent magnetic polystyrene beads (MagPlex Microspheres, Luminex) and incubated with diluted sera samples (1:50 dilution for AD-1 and AD-5/Domain I, 1:2000 for AD-4/Domain II and AD-4+5/Domain I+II). Antibody binding was detected using PE-conjugated goat-anti-mouse IgG secondary antibody (2 μg/mL, Southern Biotech). Results were acquired on a Bio-Plex 200 system (Bio-Rad) and reported as median fluorescence intensity (MFI). Binding to gB AD-2 and AD-6 was measured by 384 well plate-based ELISA at starting plasma dilution of 1:10 using methods described in previous sections and reported as AUC. The average + 3SD of baseline (Week 0) samples was used as the positivity cutoff.

### Antibody-dependent cellular phagocytosis

An optimized amount of AD169r virions (0.25 x10^3^ PFU/well) were conjugated to AF647 NHS ester prior to incubation with diluted sera samples (1:5). Then, virus-antibody immune complexes were centrifuged with 50,000 THP-1 cells for 1 hour at 1200 ×g and incubated at 37 °C for one hour. Cells were stained with Aqua Live/Dead stain, fixed with 10% formalin, and washed prior to acquisition on the flow cytometer (Fortessa; BD) using the HTS as previously described ^37,38^. The percentage of AF647+ cells was reported for each sample based on the live, singlet population. The average + 3SD of baseline (Week 0) samples was used as the positivity cutoff.

### Inhibition of cell-associated virus spread

ARPE-19 cells were infected with Ts15nR HCMV (MOI=0.05) overnight in 384-well plates. Virus was removed after 24 hours and replaced with media-containing either plasma dilutions (1:10 starting dilution),Cytogam, or media alone. Cells were stained for IE-1 expression and DAPI 12 days post infection and images were acquired and analyzed using the ImageExpress Pico Automated Cell Imaging Platform as previously described (Molecular Devices). Data was reported as the reduction in virus spread, measured by % of virus infected cells, relative to the virus control.

### Statistical Analysis and Figures

All samples were run in duplicate for all assays. Statistical analyses were performed using GraphPad Prism version 10.1.0 (GraphPad Software, Inc, La Jolla, CA) and R version 4.4.1. The statistical analyses used included Spearman rank correlation test, a paired Wilcoxon test, or Mann-Whitney U test, where appropriate; *P* < 0.05 was considered significant. P-values are adjusted for multiple comparisons using the Bonferroni method which controls for the false discovery rate (FDR). In the FDR method, p values are ranked in an ascending array and multiplied by m/k where k is the position of a p value in the sorted vector and m is the number of independent tests. Whenever appropriate, all values below the limit of detection were set to the limit of detection for statistical analyses. All graphs were generated using Graphpad Prism and figures were compiled using Adobe Photoshop. Figures 1a was generated using Biorender.

**Figure 1:**
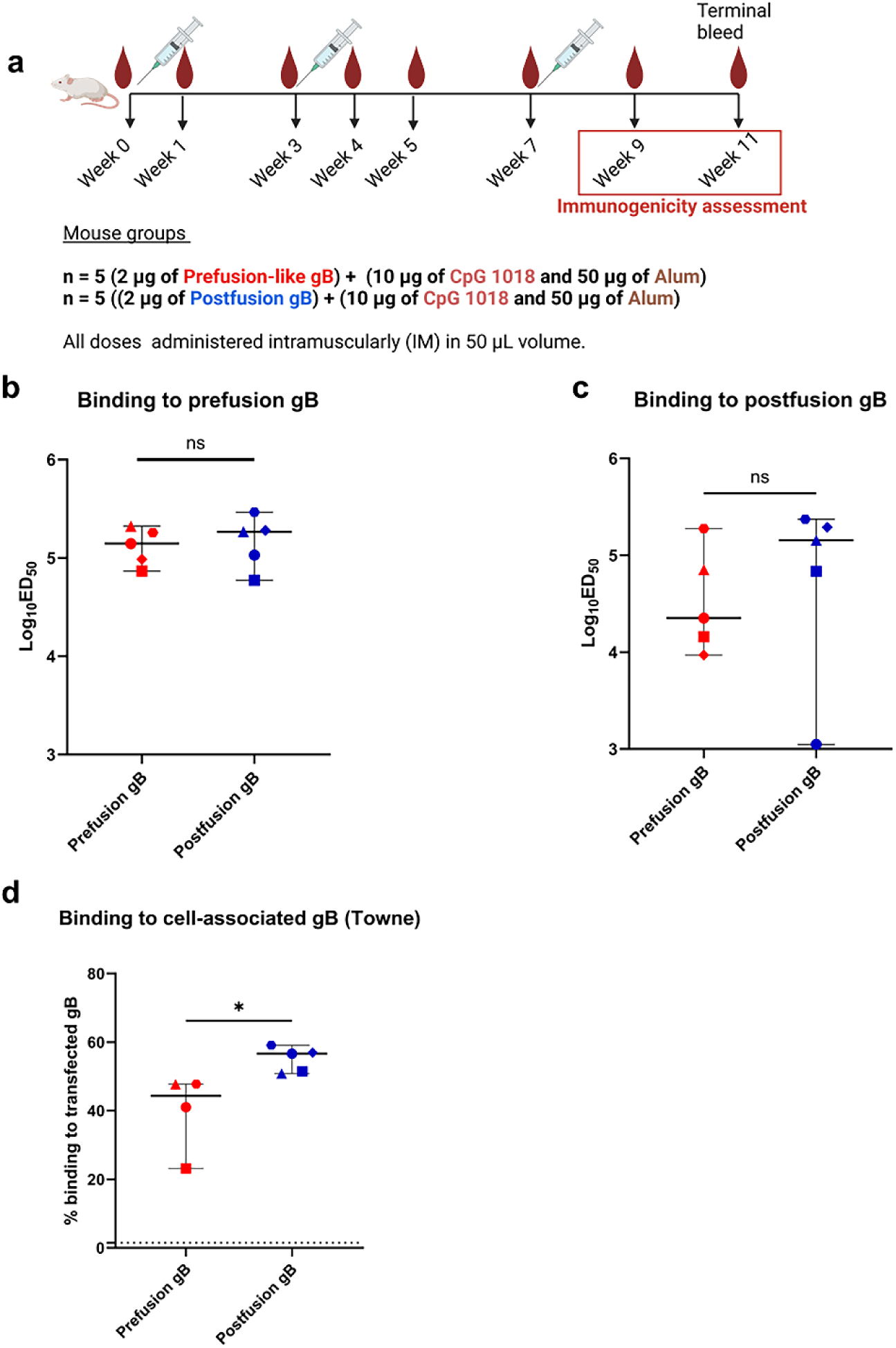
Prefusion-like gB elicited strong antibody binding to soluble and cell-associated gB. (a) 6 – 8 week BALB/c mice were immunized thrice via IM either with 2 µg prefusion-like gB or 2 µg postfusion gB, adjuvanted with CpG 1018 adjuvant and Alum. Blood was processed for plasma and immunogenicity assessments were done post all three doses (Weeks 9 and 11). Figure was generated using Biorender. Plasma IgG binding to (b) Prefusion-like gB-C7, and (c) Postfusion gB was assessed at week 9 using an ELISA and reported as Log_10_ED_50_. (d) Binding of IgG elicited by gB immunization to cell-associated Towne gB was measured by a transfected cell binding assay and reported as %binding to transfected gB. The dotted line represents the positivity cut-off, which is the average + 3 standard deviation (SD) of baseline mouse plasma. *p value < 0.05, ns = non-significant. All comparisons were done using Mann-Whitney U test in R or Wilcoxon paired t test in Graphpad Prism. Symbols denote individual mice.

## RESULTS

### Prefusion-like gB-C7 elicits strong IgG binding to soluble and cell-associated gB

Immunization with prefusion-like gB-C7 in a three-dose series (Figure 1a) elicited plasma IgG binding to prefusion-like and postfusion gB at levels comparable to that elicited by postfusion gB immunization (Figure 1b-c) at weeks 9 and 11. Prefusion-like and postfusion gB-specific IgG titers remained consistent between weeks 9 and 11 (Figure S1a). Mice immunized with prefusion-like gB-C7 demonstrated higher IgG binding to prefusion-like gB-C7 compared to postfusion gB, though this was not significantly different. Postfusion gB immunized mice elicited comparable IgG binding levels to both prefusion-like and postfusion gB (Figure 1a-b, S1b). Plasma IgG binding to cell-associated vaccine antigen-matched strain Towne was elicited by both vaccine groups. Cell-associated gB IgG binding was significantly higher in postfusion gB-immunized mice (median = 56.7%, range: 50.85 – 59.10%) compared to prefusion-like gB-C7 immunized mice (median = 44.4%, range: 23.15 – 47.75%) at week 9 (Figure 1d, p = 0.03, FDR-adjusted p = 0.06). Overall, immunization with prefusion-like and postfusion gB elicited robust IgG binding to soluble gB at comparable levels, while postfusion gB elicited higher binding to cell-associated gB compared to prefusion-like gB-C7.

### Prefusion-like gB-C7 elicited weak plasma neutralization responses at levels comparable to postfusion gB in the presence of complement

Immunization with prefusion-like and postfusion gB elicited very weak plasma neutralization of AD169r in fibroblasts by week 9. Fibroblast neutralization titers against AD169r in both vaccine groups was very low without the addition of complement, with the majority of mice from both groups demonstrating titers only slightly elevated above baseline (Log_10_ID_50_ = 1)(Figure 2a). Neutralization responses elicited by both vaccine groups were significantly increased in the presence of rabbit complement (Figure S2a-b, p = 0.008, FDR-adjusted p = 0.048 for prefusion-like gB-C7, p = 0.012, FDR-adjusted p = 0.048 for postfusion gB). Neutralization titers were weak overall and comparable between prefusion-like (median Log_10_ID_50_ = 1.25 range: 1.14 – 1.59) and postfusion gB (median Log_10_ID_50_ = 1.33, range: 1.31 – 1.50) immunized animals (Figure 2a-b), Surprisingly, neither prefusion-like nor postfusion gB immunization elicited detectable neutralization titers against Towne, the vaccine strain, at week 11 (Figure 2c), with only one of five prefusion-like gB-C7 immunized mice and no postfusion gB mice eliciting titers above that of pre-vaccine timepoints. Epithelial neutralization of AD169r in the presence of complement was nearly undetectable in all immunized mice at week 9 (Figure 2d). Overall prefusion-like and postfusion gB-elicited comparably weak and complement-dependent fibroblast neutralization of the heterologous strain AD169r, but not the vaccine strain Towne; such differences may be related to susceptibility of the virus strains towards neutralization. To explore whether human sera contain neutralizing IgG with specificities for prefusion-like or postfusion gB, we tested the neutralizing potency of Cytogam after pre-incubation with prefusion-like or postfusion gB. When pre-incubated with 3 µg or 10 µg prefusion-like gB-C7 or postfusion gB, the neutralization potency of Cytogam against AD169r in fibroblasts did not change significantly, suggesting that there is not a significant prefusion gB-specific neutralizing antibody population in HCMV hyperimmunoglobulin that is functionally distinct from that which binds to postfusion gB (Figure S3a-c).

**Figure 2:**
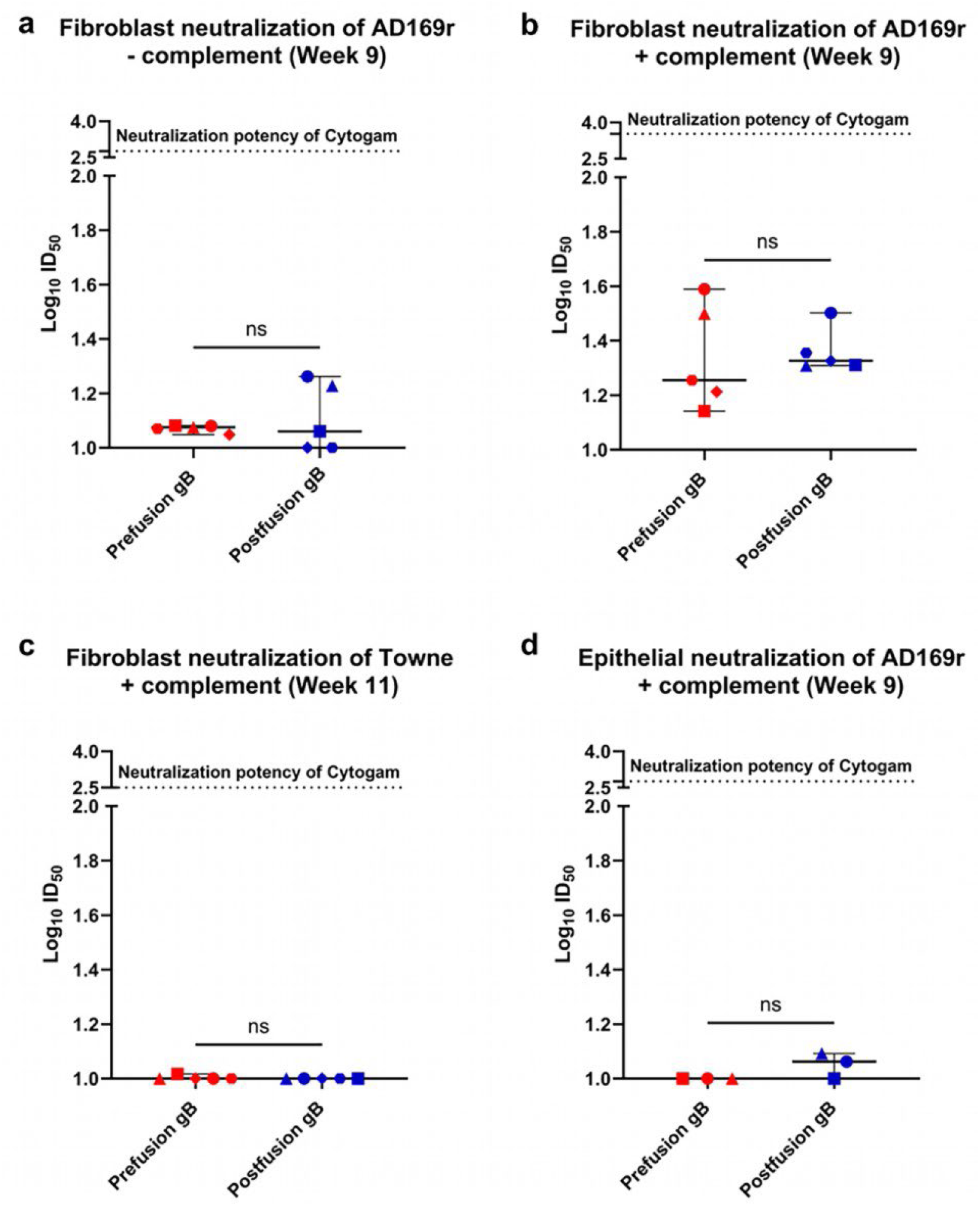
Prefusion-like gB-C7 elicited weak plasma neutralization responses at levels comparable to postfusion gB in the presence of complement. Fibroblast (HFF-1) neutralization of AD169r (a) without complement (b) with the addition of rabbit complement (1:8) was measured using a neutralization assay at week 9 and reported as Log_10_ID_50_. (c) Fibroblast neutralization of Towne at week 11 and (d) epithelial (ARPE-19) neutralization of AD169r at week 9, both with the addition of rabbit complement (1:8) were assessed using a neutralization assay and reported as Log_10_ID_50._ Dotted lines in each panel denote the neutralization potency of the hyperimmuneglobulin Cytogam. ns = non-significant. All comparisons were done using Mann-Whitney U test in R. Symbols denote individual mice.

### Postfusion gB elicited higher magnitude IgG responses to AD-5 compared to prefusion-like gB-C7

Given the lack of strong neutralizing IgG observed post vaccination despite strong binding IgG responses, we mapped vaccine-vaccination elicited IgG at week 9 to the different antigenic domains of gB (AD-1 – 6) to determine whether vaccination primarily elicited IgG against non-neutralizing domains of gB (Figure 3a). While plasma IgG binding at Week 9 to all antigenic domains was statistically higher than binding at baseline (Figure S4a-f), IgG binding was consistently above the positivity cut-off only for AD-4 (Domain II), AD-5 (Domain I), AD-4+5 (Domain I+II), and the recently characterized AD-6 (Domain V) that has been described as a target of antibodies inhibiting cell-cell spread of HCMV (Figure 3b – d)^21^. Plasma IgG elicited by both prefusion-like and postfusion gB were of highest magnitude against AD-4 on Domain II of gB (Figure 3c, measured at a dilution of 1:2000) a target of neutralizing antibodies. However, vaccine-elicited plasma IgG binding to AD-5 (Domain I) (Figure 3b, measured at a dilution of 1:50) and to the region spanning AD-4+5 (Figure 3c, measured at a dilution of 1:2000) was marginally higher for the postfusion gB group compared to prefusion-like gB-C7. While binding to AD-6 was generally low, certain mice from each group demonstrated heightened IgG binding to AD-6 (Figure 3d, measured at a starting dilution of 1:10). Surprisingly, neither prefusion-like nor postfusion gB immunization elicited detectable IgG binding to AD-1 (Figure 3b, measured at a dilution of 1:50), which has been previously described as an immunodominant AD. As expected, elicited IgG binding to AD-2 site 1, a weakly immunogenic region of gB and the target of potently neutralizing antibodies^39^, by both vaccines was extremely low (Figure 3d, measured at a starting dilution of 1:10). Additionally, we assessed correlations between vaccine-induced plasma IgG binding to the six antigenic domains to fibroblast neutralization of AD169r in the presence of complement for prefusion-like gB-C7 and postfusion gB immunization (Figure S5a-l). While the study was underpowered to achieve statistical significance, postfusion gB immunization depicted a potential trend towards a positive correlation of complement-enhanced neutralization with IgG binding to AD-6 (Figure S5l, r = 0.8, p = 0.13, Adjusted p = 0.8). Overall, vaccine-elicited IgG bound predominantly to AD-4 and AD-5, and there were statistically significant differences in IgG binding between the vaccine groups only for binding to AD-5 and AD-4+5, with postfusion gB immunization eliciting statistically higher binding to both regions (Figure 3b – c, p = 0.02, FDR-adjusted p = 0.06 for both AD-5 and AD-4+5).

**Figure 3:**
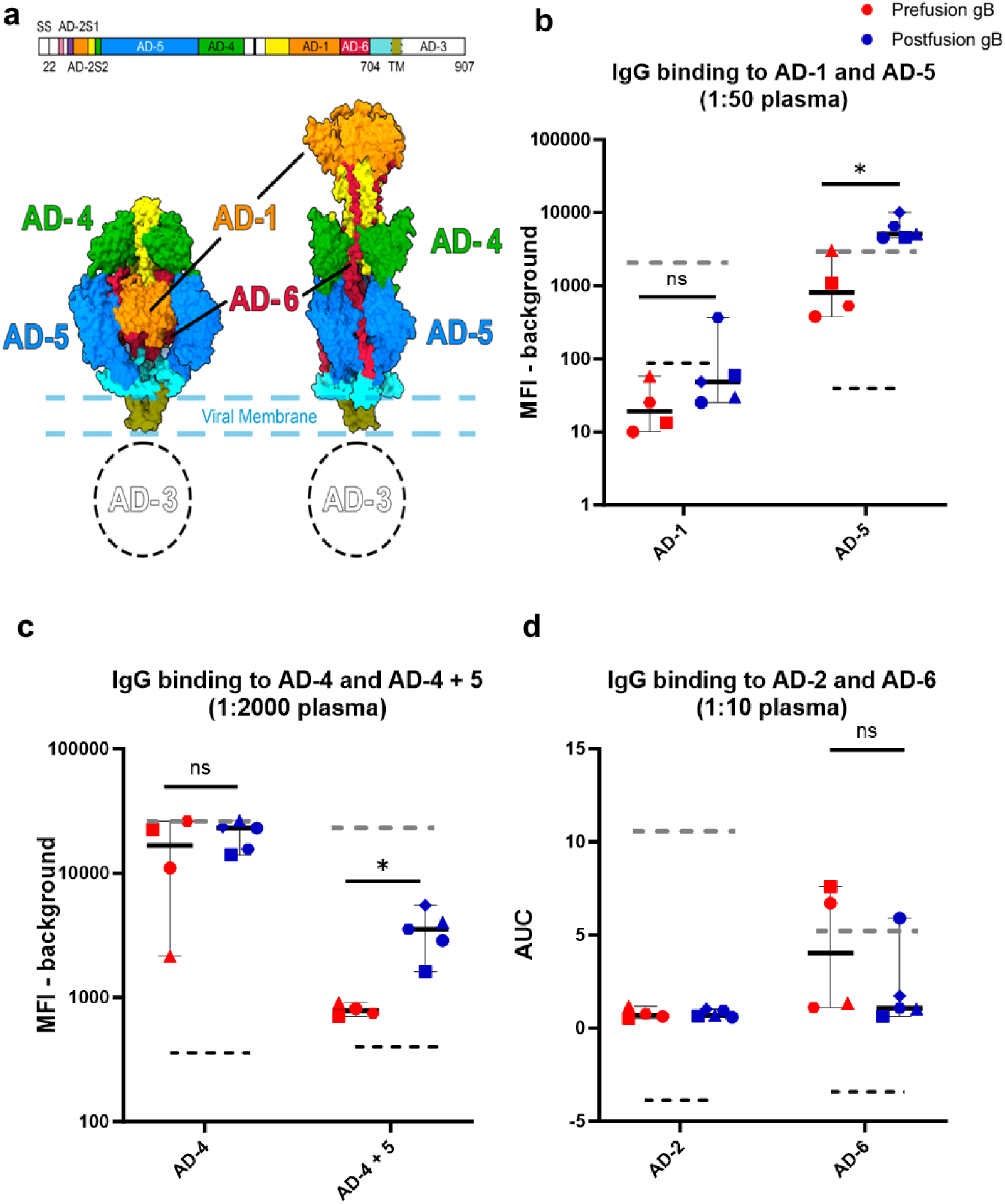
Vaccine-elicited IgG mapped to the neutralizing epitope AD-4 and AD-5 on gB, with significantly higher AD-5 binding induced by postfusion gB. (a) Linear and structural schematic depicting the location of the six antigenic domains (AD) of gB in the prefusion and postfusion state. Plasma IgG binding elicited by prefusion-like and postfusion gB was assessed to (b) AD-1 and AD-5, at a 1:50 plasma dilution, and to (c) AD-4 and AD-4+5 at a 1:2000 plasma dilution using BAMA and reported as background subtracted median fluorescence intensity (MFI). (e) Vaccine induced IgG binding to AD-2 and AD-6 at 1:10 plasma dilution was measured using ELISA and reported as Area under the Curve (AUC). Black dotted lines represent the positivity cut-off, which is the average + 3 standard deviation (SD) of baseline mouse plasma. Gray dotted lines represent binding of the positive control Cytogam to respective ADs. *p value < 0.05, ns = non-significant. All comparisons were done using Mann-Whitney U test in R. Symbols denote individual mice.

### Prefusion-like gB-C7 immunization elicited lower levels of ADCP relative to postfusion gB

As the importance of non-neutralizing antibodies in conferring protection against HCMV acquisition and congenital transmission has been demonstrated in several studies^37,40,41^, we assessed ADCP responses against the HCMV AD169r strain at week 9 post vaccination in comparison to unmatched seronegative (baseline) mouse plasma. ADCP was measured using THP-1 monocytes at a single plasma dilution (1: 5) and reported as % AF647 as a measure of phagocytosis of AF647-conjugated AD169r. Prefusion-like and postfusion gB immunization elicited strong ADCP responses compared to baseline (Figure 4a), but postfusion-gB immunized mice elicited significantly higher plasma ADCP responses (median = 21.5%, range 15.75 – 23%) compared to prefusion-like gB-C7 immunization (median = 14.1%, range 13 – 18.55%) (Figure 4a, p = 0.03, FDR-adjusted p = 0.03).

**Figure 4:**
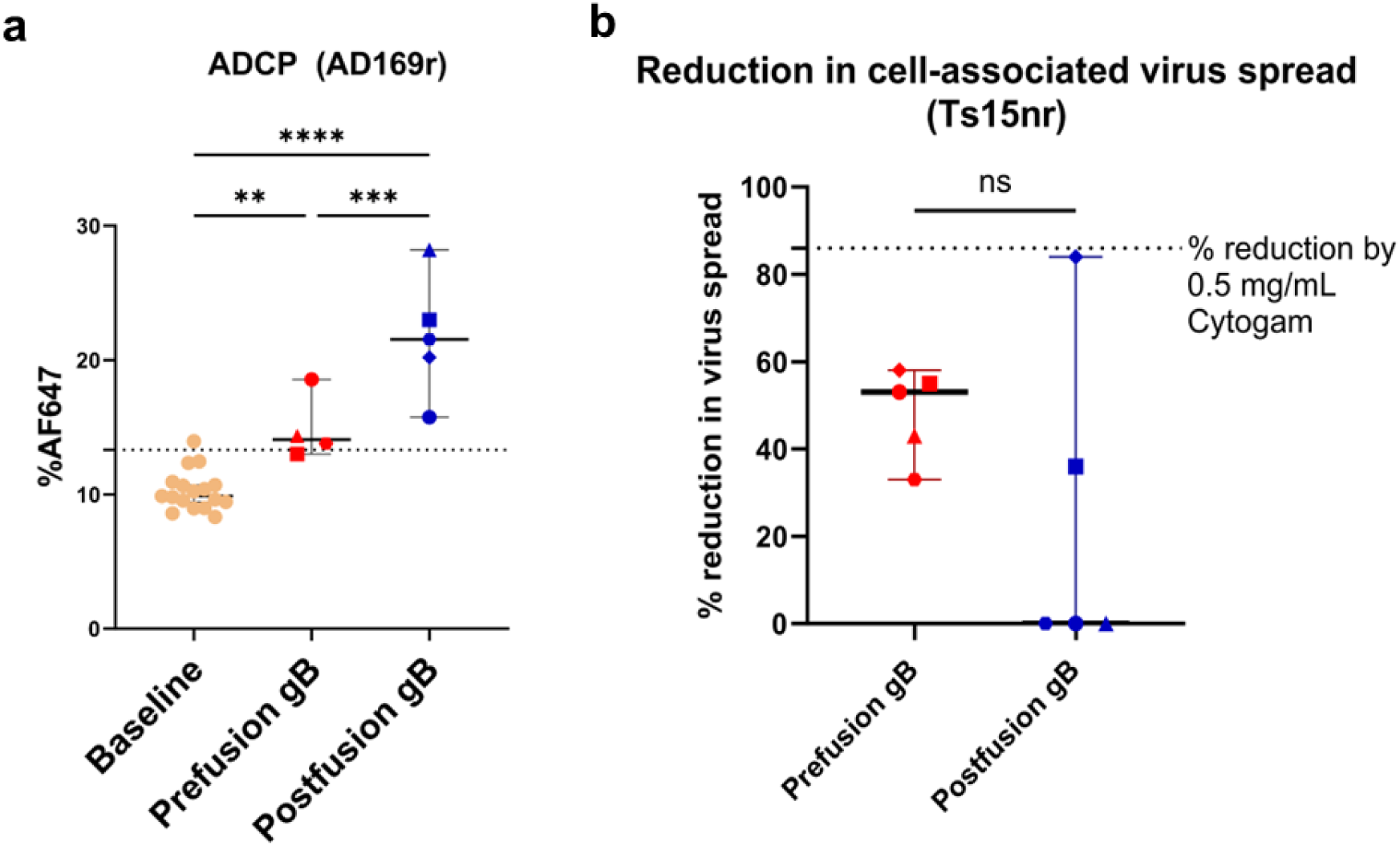
Prefusion-like gB-C7 immunization elicits non-neutralizing antibody responses that exhibit lower ADCP compared to postfusion gB immunization. (a) Baseline (unpaired mouse plasma) and vaccine-elicited ADCP responses against AD169r was assessed in THP-1 monocytic cell line and reported as %AF647. The dotted line represents the positivity cut-off, which is the average + 3 standard deviation (SD) of baseline mouse plasma. (B) Reduction in cell-associated spread of Ts15nr was assessed at 1:10 plasma dilution and reported as %reduction. The dotted line represents reduction induced by 0.5 mg/mL Cytogam. **p<0.01, ***p<0.001, ****p<0.0001, ns = non-significant. All comparisons were done using Mann-Whitney U test in R or one-way ANOVA in Graphpad Prism. Symbols denote individual mice.

### Prefusion-like gB-C7 immunization elicits antibodies that reduce cell-associated virus spread

As half of the mice immunized with prefusion-like gB-C7 elicited IgG responses against AD-6, a domain shown to elicit non-neutralizing antibodies implicated in cell-cell spread of HCMV, we were interested in evaluating the extent to which each vaccinated group could inhibit cell-cell spread of HCMV. We tested the ability of plasma IgG at week 11 to inhibit cell-associated virus spread of HCMV strain Ts15-nR, a BAC variant of Towne, the vaccine strain, with a pentameric reversion that restores epithelial cell tropism. Plasma IgG elicited by prefusion-like gB-C7 exhibited modest inhibition of cell-associated spread of Ts15nr relative to average spread observed in control conditions (Median reduction = 53%, range 33 – 58%) (Figure 4b, p and Adjusted p value = 0.205). Postfusion gB immunization elicited lower magnitude inhibition in virus spread compared to prefusion-like gB-C7-immunized mice, although this difference was not statistically significant (median reduction = 0%, range 0 – 84%). Only 2 out of 5 postfusion gB immunized mice demonstrated plasma IgG that blocked cell-associated virus spread, while all 5 out of 5 prefusion-gB immunized mice mounted IgG that could inhibit virus spread. We also assessed the correlation between reduction in cell-associated virus spread and IgG binding to AD-6 for both prefusion-like gB-C7 (Figure S6a) and postfusion gB (Figure S6b). Though the study was statistically underpowered to achieve significance, Spearman’s correlation indicated a trend towards a positive correlation between prefusion-like gB-elicited reduction in cell-cell virus spread and IgG binding to AD-6, with a p value that approached statistical significance (Figure S6a r= 1, p = 0.08, Adjusted p = 0.17). Overall, prefusion-like gB-C7 immunization induced plasma IgG that can block cell-associated virus spread of Ts15nR, which may be mediated by AD-6 specific IgG.

## DISCUSSION

The successful development of a vaccine against HCMV has proved to be quite challenging despite concerted efforts over five decades, due in part to a need to better understand the immune correlates of protection and conformational requirements for an efficacious vaccine. While most prior vaccine efforts focused on gB as a key glycoprotein antigen, gB in its postfusion state has typically elicited low neutralizing antibody titers^40^. Recent vaccine candidates have focused on other glycoproteins such as PC^42,43^, or employed unique delivery platforms for HCMV glycoproteins, such as the enveloped virus-like particle (eVLP) system used to deliver membrane-bound gB^13^. An HCMV-Modified Vaccinia Ankara (MVA) triplex vaccine encoding pp65, IE1, and IE2 is in Phase 2 trials^44^, while the mRNA-1647 vaccine (Moderna) encoding gB and PC has progressed to Phase III trials after eliciting strong neutralizing antibody and cellular immune profiles in Phase II participants^42,43^. Class I viral fusion proteins stabilized in the prefusion conformation have elicited robust neutralizing antibody titers at higher magnitude compared to their postfusion conformations.^45,46^ These efforts have culminated in the FDA-approval of prefusion-stabilized vaccines using class I viral fusion proteins in vaccines against SARS-CoV-2 and RSV. Inspired by these successes, there has been interest in stabilizing the prefusion conformation of HCMV gB, a class III fusion protein, and test its potential as a vaccine candidate to improve virus neutralization responses. However, efforts were stymied by the lack of any prefusion structures for herpesvirus gB homologs. When a structure of detergent-solubilized full-length gB structure maintained in the prefusion conformation by a fusion inhibitor and chemical crosslinking was published in 2021, this enabled structure-based design efforts for gB in its metastable prefusion state^34^. Recently, a soluble HCMV gB ectodomain which used amino acid substitutions to stabilize gB in a prefusion-like conformation, exhibited improved expression and thermostability relative to soluble postfusion gB ectodomain^35^. However, this HCMV gB prefusion-like construct, referred to as gB-C7, did not able to elicit higher fibroblast neutralizing antibody responses despite inducing higher IgG binding titers post vaccination^35^ (Figures 1b-c, 2a-d). In this study, we further characterized the humoral immune response to the recently described prefusion-like HCMV gB construct gB-C7 relative to postfusion HCMV gB.

The weak neutralizing responses elicited by subunit gB vaccination have been observed previously, most notably in the gB/MF59 vaccine trial^14,15,40^. In agreement with the initial immunization results published on HCMV gB-C7, our results showing that gB stabilized in a prefusion-like conformation could not improve upon these poor neutralizing titers. This suggests that unlike class I prefusion-stabilized immunogens, prefusion-like gB-C7 does not display neutralization-sensitive epitopes to a better extent than postfusion gB^35^. Surprisingly, while both vaccine groups elicited weak, complement-mediated fibroblast neutralization of the heterologous, lab-adapted HCMV strain AD169r, neither group exhibited any detectable neutralization of the vaccine-matched strain Towne. This result is in direct contradiction to the gB/MF59 vaccine study, where complement-mediated neutralization of the vaccine strain Towne was higher than that observed against AD169r in the adolescent population^40^. While these could be attributed to virologic differences between AD169r and Towne^47^ and the hypervariable nature of gB between virus strains, it is important to evaluate the specificity of the weak neutralization exhibited against AD169r in the context of protection against HCMV acquisition or transmission, which is beyond the scope of this study. There was also no detectable neutralization of AD169r in epithelial cells observed in the presence of complement, which is consistent with lack of responses seen in phase II postpartum participants in the gB/MF59 vaccine trial^40^.

The weak neutralization responses are surprising in the context of the predominant AD-4 IgG response against both gB constructs evaluated in this study, which has been a target of broadly neutralizing anti-HCMV antibodies^31^. Furthermore, while both conformations of gB display AD-5 on Domain I (Figure 3a), in our study, postfusion gB immunization elicited significantly higher IgG binding to AD-5 and the AD-4+5 region (Figure 3b-c), which could indicate that immunodominant epitopes on AD-5 are better presented in the postfusion structure. Neither prefusion-like nor postfusion gB immunization was able to elicit IgG titers against AD-2 site 1, a weakly immunogenic target of potently neutralizing antibodies such as TRL345^39^. AD-2 site 1 is a highly conserved region in the N terminus of gB, and its structure has not been resolved in either prefusion or postfusion states, suggesting it may be intrinsically flexible. In fact, no vaccine has elicited consistent binding or neutralizing antibody responses against the AD-2 site 1 region, though AD-2 responses are reported in approximately half of seropositive individuals^48^ and have been correlated with decreased post-transplant viremia^49^. Structures of prefusion or postfusion gB that can resolve the AD-2 site 1 region could lead to the design of a gB construct that may better elicits AD-2 site 1-specific IgG and enhance the neutralizing antibody responses against gB. Thus, future studies of prefusion gB designs should prioritize display of this region and evaluate how it interacts with germline B cell lineages that have naturally resulted in potent AD-2 site1-specific broadly neutralizing antibodies^50^. While future iterations of structure-guided design of prefusion gB could yield better engagement of broad and potent neutralizing antibody precursors, the extensive glycosylation of gB could present barriers to eliciting neutralizing antibody responses and could be addressed through further antigen design^51^.

While neutralization has traditionally been regarded as the key immunologic target of HCMV vaccines, it has become clear than neutralization alone is likely insufficient in mediating protection against transmission. The importance of non-neutralizing antibody functions for protection against HCMV transmission has been demonstrated especially in the setting of congenital HCMV transmission. This involves the engagement of FcγRI and FcγRIIA as well as Fc-mediated effector functions such as antibody dependent cellular phagocytosis (ADCP) and antibody dependent cellular cytotoxicity (ADCC) contributing to reduced transmission in plasma and cord blood of transmitting and non-transmitting mother-infant dyads^37,41^. Our group has also shown that the partial protection observed in the Phase IIb gB/MF59 cohort was potentially due to Fc-mediated effector functions such as virion phagocytosis rather than the weakly-elicited neutralizing responses^40^. Another recent study associated this protection to inhibition of HCMV spread in infected cells, mediated through the recently identified non-neutralizing AD-6 domain of gB^21^. In this mouse immunogenicity study, postfusion gB elicited significantly higher ADCP against AD169r compared to immunization with prefusion-like gB-C7, one of the main differences identified in the immunogenicity between these two conformations. This corroborates the high ADCP observed with the gB/MF59 vaccine^40^, significantly higher than that elicited by the mRNA-1647 gB+PC vaccine^43^. Further characterization of the distinct gB epitopes and gB-specific IgG engagement with Fcγ receptors elicited by these fusion states could shed light on the observed differences in ADCP. On the other hand, prefusion-like gB-C7 immunization elicited IgG that inhibited epithelial cell-associated virus spread of Ts15nR at higher levels and in a higher proportion of animals than observed after postfusion gB immunization, though this difference was not statistically significant. Further, most animals with detectable AD-6-specific IgG also displayed reduction in epithelial cell spread of Ts15nR.

HCMV vaccine studies have also focused on cell-associated gB assayed through transfected or infected cell binding as an important immunogenicity measure, as IgG binding to gB on the surface of a cell was identified as a correlate of protection in seronegative women in the gB/MF59 vaccine trial^19^. While native gB on the infected cell surface is present in both the metastable prefusion and the stable postfusion conformations^34^, the prefusion state is more common. Interaction with the gH/gL complex is thought to the provoke the conformational rearrangement of gB to the stable postfusion form, a process which mediates fusion between the viral and host cell membrane for viral entry^26^. Though both vaccine groups elicited high titers of IgG against cell-associated gB, it is possible that the higher binding to cell-associated gB observed with postfusion gB immunization could be due to the lack of conformationally-specific epitopes on prefusion-like gB-C7. Prefusion-like gB-C7 did not preferentially block Cytogam-mediated neutralization differently than postfusion gB (Figure S4). This suggests that natural HCMV infection elicits neutralizing antibodies that may not be specific to these particular gB conformations, supported by the lack of any published reports on prefusion-gB specific antibodies to date and the limited neutralizing antibodies elicited by these vaccines. Additionally, it is possible that prefusion-like gB-C7 elicits reduced IgG binding to cell-associated gB due to structural differences between the soluble prefusion-like gB-C7 ectodomain evaluated here and the native prefusion conformation of full-length membrane-bound HCMV gB expected to predominate on the surface of a cell.

Our murine immunogenicity analysis explores the lack of enhancement in neutralizing responses observed post vaccination with prefusion-like gB-C7 in mice and characterizes several non-neutralizing IgG functions as well as binding to cell-associated gB (Supplementary Table 1), the small sample size limits further exploratory analysis. Overall, prefusion-like gB-C7 does not appear to be more immunogenic for functional anti-viral antibody responses compared to the postfusion conformation. Future iterations of engineered gB constructs that optimize the display of key epitopes could result in improved immunogenicity profiles. HCMV vaccines that present optimized HCMV glycoproteins and utilize novel delivery platforms to elicit responses associated with protection against HCMV disease and congenital transmission could be key to developing successful HCMV vaccine candidates.

## DATA AVAILABILITY

All datasets generated and analyzed during the current study are available in the main text and supplementary information of this manuscript. Datasets used for figure generation and statistical analysis will also be made available on Github upon publication.

## ACKNOWLEDGEMENTS

This study was funded by Dynavax Technologies Corporation.

## AUTHOR CONTRIBUTIONS

S.R.P. oversaw and conceptualized the project. S.R.P. and K.P.K designed the study. K.P.K. planned the animal studies and processing while C.R.B., M.G., C.A.W., and I.M. performed animal procedures (handling, immunization, sample collection, processing). K.P.K led the optimization and performance of the immunoassays as well as the data processing, figure generation, and analysis. M.C. performed and analyzed binding, neutralization, and cell spread immunoassays and helped in manuscript preparation. A.S.F., C.M.C., and S.H. designed, optimized, and performed the antigen domain mapping and data analysis. Antigens were designed and provided by M.R.S., P.O.B., and J.S.M. The CpG1018 adjuvant was provided by M.B. C.A. and L.G. performed statistical analysis. K.P.K. drafted the paper with editing from co-authors, and all authors reviewed the paper draft before submission.

