## Supplementary Material for "A human cytomegalovirus prefusion-like glycoprotein B subunit vaccine elicits similar humoral immunity to that of postfusion gB in mice"

**Supplementary Figures**

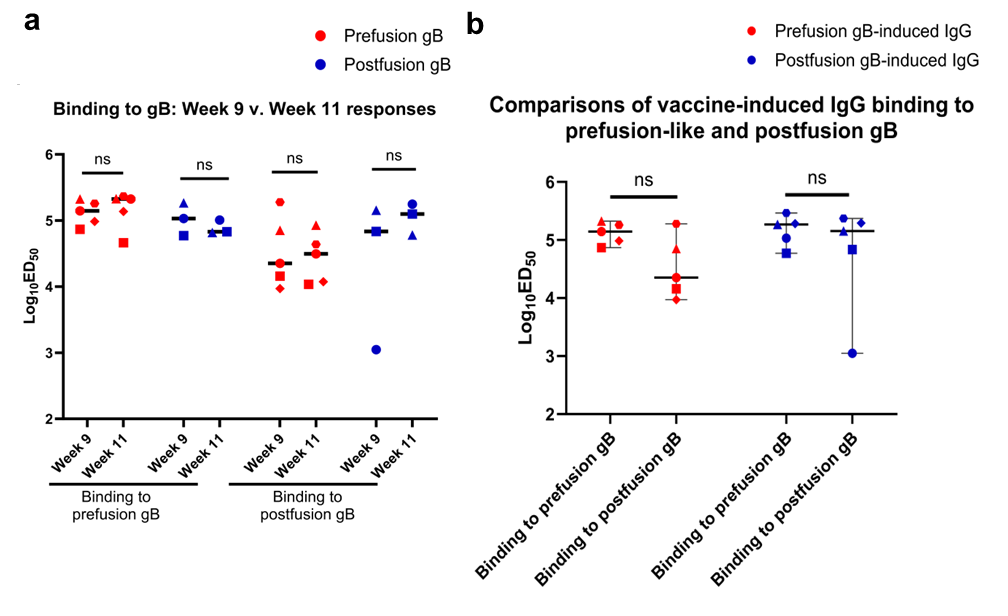
**Figure S1: Vaccine-induced IgG binding to soluble and cell-associated gB** (a) Prefusion and postfusion gB immunization-induced IgG binding to soluble antigens prefusion and postfusion gB was assessed using an ELISA and reported as Log_10_ED_50_. (b) Plasma IgG binding to Prefusion-like gB-C7, and Postfusion gB was compared at weeks 9 and 11 using an ELISA and reported as Log_10_ED_50._ Symbols denote individual mice. All comparisons were done using Wilcoxin ranked sum test in Graphpad Prism.

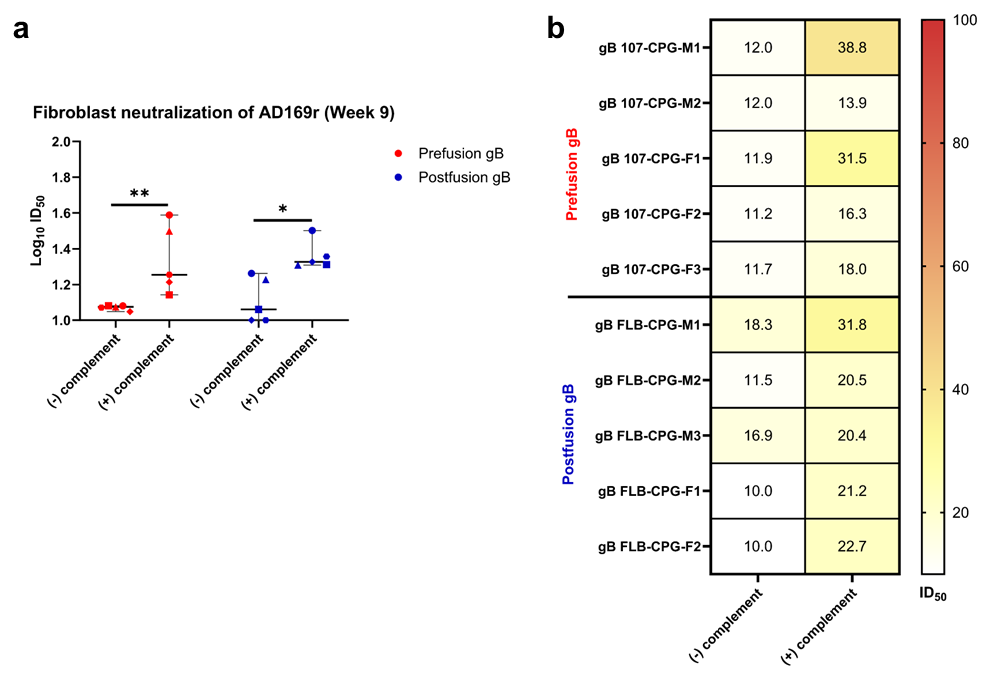
**Figure S2: Complement enhances fibroblast neutralization of AD169r elicited by prefusion-like and postfusion gB vaccination.** (a) Fibroblast neutralization of AD169r was assessed without and with the addition of rabbit complement (1:8) in both prefusion and postfusion gB vaccine groups and reported as Log_10_ID_50_ . (b) Heat map representing the ID_50_ of fibroblast neutralization of AD169r assessed in plasma of individual mice without (-) and with (+) complement. *p<0.05, **p<0.01. All comparisons were done using Wilcoxin ranked sum test in R.

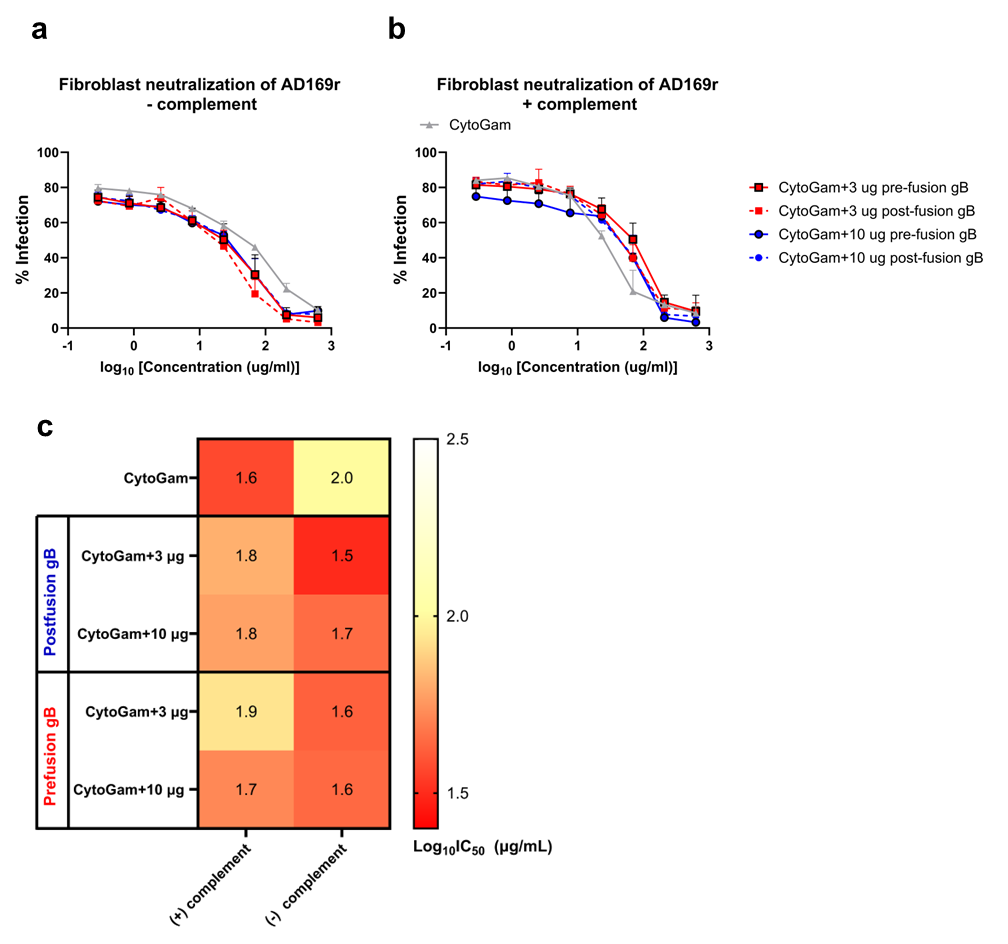
**Figure S3: Prefusion or postfusion gB do not block neutralizing activity of cytogam.** Fibroblast neutralization of AD169r elicited by Cytogam was assessed alone or in the presence of 3 µg (dotted line) or 10 µg (solid line) prefusion or postfusion gB (a) without complement and (b) with the addition of rabbit complement. (c) Heat map depicting the Log_10_IC_50_ of Cytogam-induced neutralization of AD169r when blocked with prefusion or postfusion gB antigens with and without complement is shown.

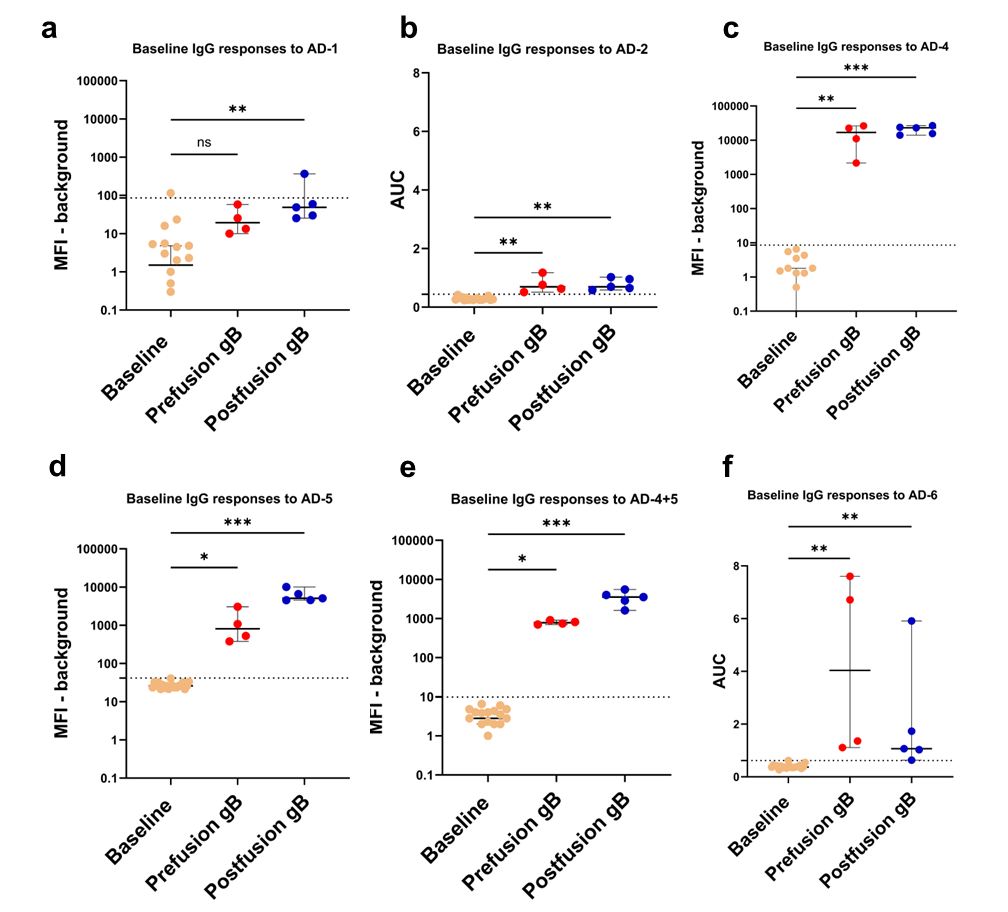
**Figure S4: Baseline mouse IgG binding to gB antigenic domains.** Baseline IgG binding to (a) AD-1 (b) AD-2 (c) AD-4 (d) AD-5 (e) AD-4+5, and (f) AD-6 was assessed in unpaired mouse plasma. Binding to AD-1, AD-4, AD-4+5, and AD-5 was measured using BAMA and reported as background subtracted MFI. Binding to AD-2 and AD-6 was measured using ELISA and reported as AUC. Baseline samples were used to calculate the positivity cut-off for each antigen, represented by dotted lines on the graph as average + 3 SD of baseline IgG binding. *p<0.05, **p<0.01, ***p<0.001, ns = non-significant. All comparisons were done using one-way ANOVA in Graphpad Prism.

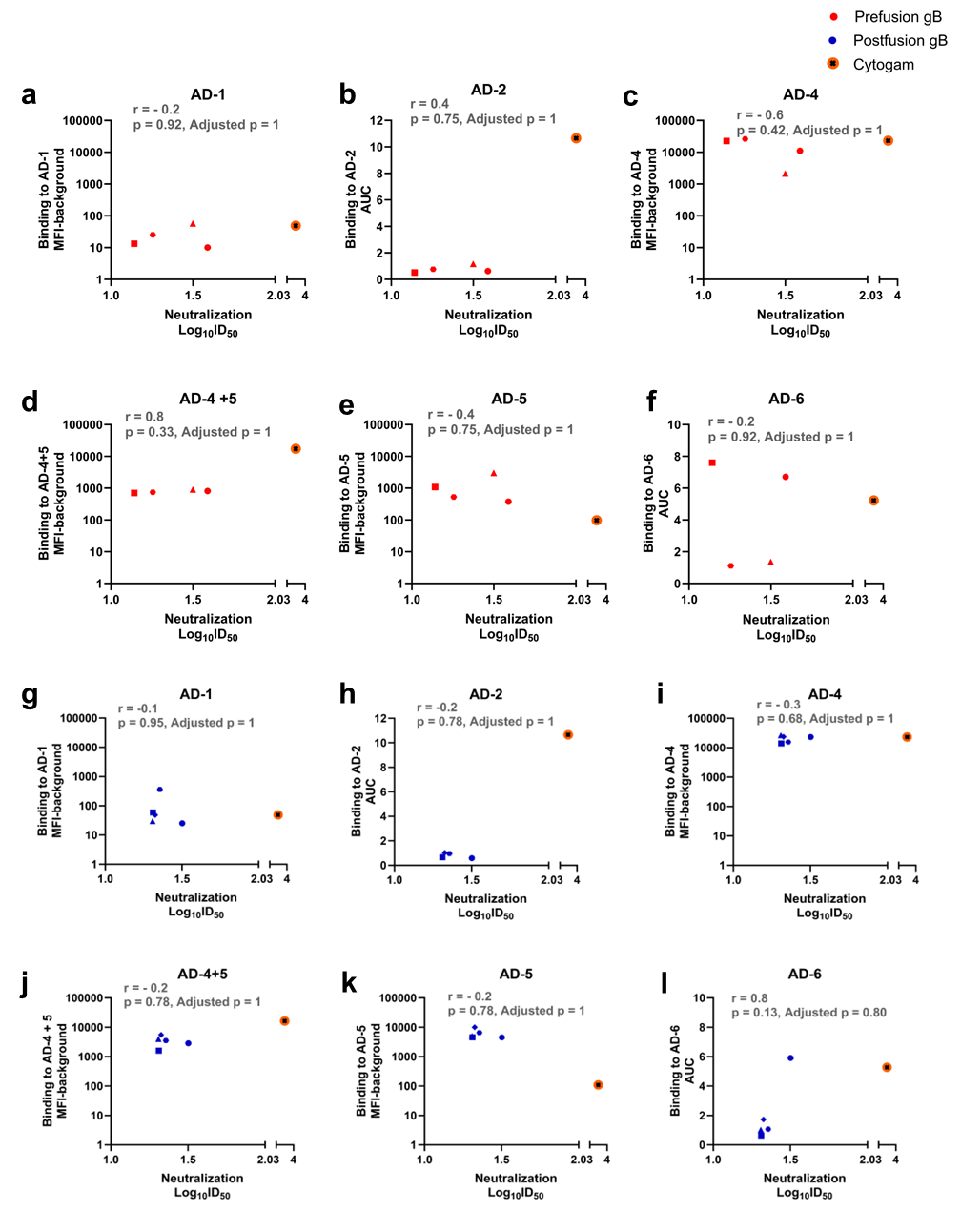

**Figure S5: Correlation between IgG binding to gB antigenic domains and fibroblast neutralization of AD169r with complement elicited by prefusion-like gB-C7 and postfusion gB vaccination** Correlation between prefusion-like gB induced IgG binding to (A) AD-1, (B) AD-2, (C) AD-4, (D) AD-4+5, (E) AD-5, and (F) AD-6, postfusion gB induced IgG binding to to (G) AD-1, (H) AD-2, (I) AD-4, (J) AD-4+5, (K) AD-5, and (L) AD-6, and fibroblast neutralization of AD169r with the addition of complement induced by Prefusion gB immunization was assessed using Spearman’s non-parametric correlation test in Graphpad Prism. Insets report Spearman’s r and p values. Cytogam is plotted as a reference but was not used in correlation analysis.

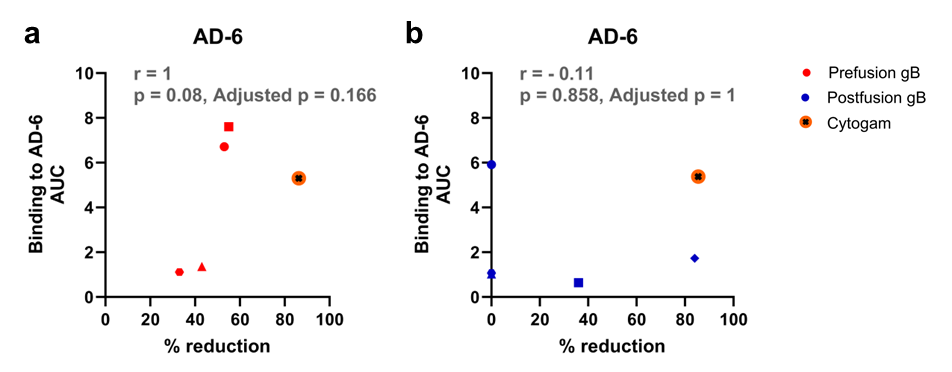

**Figure S6: Correlation between IgG binding to AD-6 and reduction of cell-associated virus spread of Ts15nr** Correlation between IgG binding to AD-6 (AUC) and %reduction of cell-associated virus spread of Ts15nr induced by (a) Prefusion gB and, (b) Postfusion gB immunization was assessed using Spearman’s non-parametric correlation test in R. Insets report Spearman’s r and p values. Cytogam is plotted as a reference but was not used in correlation analysis.

| Humoral Response | Prefusion gB immunized mice  (n = 3-5) | Postfusion gB immunized mice  (n = 3-5) | p-value | FDR-adjusted p-value |
| --- | --- | --- | --- | --- |
|  | Median (Range) | Median (Range) |  |  |
| **IgG binding** |  |  |  |  |
| *Soluble gB-specific (Log_10_ED_50_)* |  |  |  |  |
| Binding to prefusion gB |  |  |  |  |
| Week 9 | 5.146 (4.867 , 5.324) | 5.267 (4.772, 5.466) | 0.690 | 0.841 |
| Week 11 | 5.326 (4.662 , 5.368) | 4.831 (4.818 , 5.007) | 0.25 |  |
| Binding to postfusion gB |  |  |  |  |
| Week 9 | 4.352 (3.971 , 5.277) | 5.155 (3.048 , 5.373) | 0.421 | 0.841 |
| Week 11 | 4.496 (4.033 , 4.640) | 5.098 (4.782 , 5.247) | 0.071 |  |
| *Cell-associated gB (%binding)* |  |  |  |  |
| Towne | 44.38 (23.15 , 47.75) | 56.65 (50.85 , 59.10) | 0.029 | 0.056 |
| **Neutralization (Log_10_ID_50_)** |  |  |  |  |
| AD169r (fibroblast) – C | 1.075 (1.048 , 1.081) | 1.060 (1.000, 1.262) | 0.834 | 0.834 |
| AD169r (fibroblast) + C | 1.255 (1.142 , 1.589) | 1.326 (1.309 , 1.502) | 0.548 | 0.626 |
| Towne (fibroblast) | 1.000 (1.000 , 1.017) | 1.000 (1.000 , 1.000) | 0.617 | 0.617 |
| AD169r (epithelial) | 1.000 (1.000 , 1.000) | 1.062 (1.000 , 1.092) | 0.197 | 0.386 |
| **gB AD mapping (MFI)** |  |  |  |  |
| AD-1 | 19.3 (10.0 , 57.8) | 48.5 (25.3 , 365) | 0.140 | 0.201 |
| AD-2 site 1 (AUC) | 0.693 (0.513 , 1.177) | 0.693 (0.586 , 1.025) | 0.905 | 0.905 |
| AD-4 | 16717.9 (2151.3 , 26265.0) | 23049 (14056.5 , 26453.0) | 0.413 | 0.561 |
| AD-5 | 805. 1 (378.3 , 3041.5) | 5096.5 (4531.0 , 10073.0) | 0.016 | 0.063 |
| AD-4&5 | 779.5 (702.5 , 906.8) | 3534.8 (1611.3 , 5524.0) | 0.016 | 0.063 |
| AD-6 (AUC) | 4.036 (1.109 , 7.607) | 1.069 (0.633 , 5.910) | 0.190 | 0.619 |
| **Non-neutralizing antibody functions** |  |  |  |  |
| ADCP (% Phagocytosis) | 14.075 (13 , 18.55) | 21.55 (15.75 , 23) | 0.032 | 0.032 |
| % reduction in cell-associated spread | 53 (33, 58) | 0 (0, 84) | 0.205 | 0.205 |

**Supplementary Table 1: Summary of plasma humoral responses assessed in mice immunized with prefusion-like gB or postfusion gB**

MFI: Median fluorescence intensity FDR: False discovery rate
